# Axonal transport and active zone proteins regulate volume transmitting dopaminergic synapse formation

**DOI:** 10.1101/284042

**Authors:** David M. Lipton, Celine I. Maeder, Kang Shen

**Author notes:** Correspondence should be addressed to K.S.

## Abstract

At a typical synapse, the precise juxtaposition between the active zone and postsynaptic receptors ensures local and precise neurotransmitter release and detection. Dopamine neurons release neurotransmitter more diffusely using volume-transmission, where precise pre- and post-synaptic alignment is lacking. It is unknown whether Dopaminergic presynaptic terminals have typical active zone structures and how they develop. Here we show that presynaptic terminals of the *C. elegans* dopaminergic neuron PDE contain *bona fide* AZ proteins, including SYD-2/Liprin-ʡ, ELKS-1, UNC-10/RIM and CLA-1/Piccolo. During development, synaptic vesicles (SVs) and active zone proteins (AZs) coalesce within minutes behind the advancing growth cone. Precise regulation of UNC-104/Kinesin-3-mediated SV transport through kinesin autoinhibition is required to pause transported SVs at synapses. SYD-1 and SYD-2 recruit and cluster the transiting SVs, while ELKS-1 aggregates through a distinct mechanism.

## Introduction

The function of neural circuits depends on the precise assembly of synapses between appropriate partner neurons, through the process of synaptogenesis. At a typical synapse, such as a glutamatergic, cortical or hippocampal synapse, presynaptic specializations in the axon contain synaptic vesicles (SV) and active zone structures where neurotransmitter release takes place. Active Zone proteins (AZs) dock and prime SVs for efficient release, recruit calcium channels to the adjacent pre-synaptic membrane, and coordinate the positioning of pre-and post-synaptic specializations via trans-synaptic cell-adhesion molecules (Petzoldt et al., 2016; Südhof, 2012). At most synapses, the post-synaptic specialization contains neurotransmitter receptors as well as an intricate set of structural and signaling proteins that are important for synapse development, function, and plasticity.

Dopaminergic neurons release the neuro-modulator dopamine in brain regions such as striatum, and cortex, and play essential roles in important behaviors such as reward learning and addiction (Hyman et al., 2006). In contrast to classic glutamatergic synapses, dopaminergic synapses are much less well studied. One key feature of the dopamine synapse is communication by a form of neurotransmitter release known as volume transmission (Agnati et al., 2010), where dopamine is diffusely released onto target brain regions from widely arborized terminals, most of which are not directly opposed to a post-synaptic density and spine (Matsuda et al., 2009; Sulzer et al., 2016; Trudeau et al., 2014; Zhang et al., 2015). This unique anatomy of dopamine pre-synapses, in addition to the recent discovery that many of these dopamine pre-synaptic release sites are functionally silent (Pereira et al., 2016) highlights the need to further characterize these neuro-modulatory pre-synaptic release sites (Sulzer et al., 2016; Trudeau et al., 2014).

In particular, both the molecular composition of dopamine pre-synaptic terminals (Pereira et al., 2016; Trudeau et al., 2014) and the synaptogenesis events that lead to their formation are still largely unknown. 1. Do dopamine neurons contain bona fide Active Zones at their pre-synaptic terminals? 2. How are synaptic vesicles and other pre-synaptic materials transported to the developing *en passant* pre-synaptic terminals? 3. What is the precise sequence of coalescence events of both SVs and AZs, if present, which lead to the formation of synapses *in vivo*, behind an extending growth cone? These last two questions are of universal importance and apply to all types of synapses.

Synaptic vesicles are thought to arise from synaptic vesicle precursors (SVPs), which are generated in the cell body and transported down the axon by UNC-104/Kinesin-3 motors along microtubule tracks (Hall and Hedgecock, 1991; Okada et al., 1995; Pack-Chung et al., 2007) before pausing and accumulating at synaptic sites (Jin and Garner, 2008; Sabo et al., 2006). Recent studies show that the SV bound small GTPase ARL-8 activates UNC-104/Kinesin-3 to ensure the delivery of SVs to the distal axon (Klassen et al., 2010; Wu et al., 2013). ARL-8 activates UNC-104/Kinesin-3 by releasing this kinesin’s autoinhibition, a state of motor inactivation that is shared by many kinesins (Hammond et al., 2009; Niwa et al., 2016).

At the same time, a subset of Active Zone proteins (AZs), most notably SYD-1 and SYD-2, play crucial roles in recruiting vesicles to the synapse, as loss of these crucial synapse assembly promoting active zone proteins reduces the number of vesicles clustered at the synapse (Dai et al., 2006; Patel et al., 2006; Zhen and Jin, 1999) and reduces the size of active zone structures, called dense projections (Kittelmann et al., 2013). Presynaptic assembly proteins at the active zone, such as *syd-2*, promote SV aggregation, leading to the model of an antagonistic relationship between SV trafficking and AZ mediated synapse formation (Klassen et al., 2010; Maeder et al., 2014; Wu et al., 2013).

After their synthesis in the cell body (Dresbach et al., 2006; Maas et al., 2012), AZ proteins are thought to be incorporated into “Piccolo-Bassoon transport vesicles” (PTVs) (Shapira et al., 2003; Zhai et al., 2001; Ziv and Garner, 2004). These vesicles are then thought to be transported down the axon to sites of synapse assembly, where a relatively small number of mobile PTVs coalesce to form the active zone (Shapira et al., 2003). Yet, these PTV transport packets have been difficult to observe *in vivo* (Petzoldt et al., 2016). In addition, there is no consensus about which motors are used for AZ transport (Goldstein et al., 2008). In some contexts, AZs are at least partially dependent on Kinesin-3 for trafficking to the synapse (Klassen et al., 2010; Pack-Chung et al., 2007) and in others they are Kinesin-3 independent (Klassen et al., 2010; Patel et al., 2006). There is also evidence of AZs being co-trafficked along the axon with SVs *in vitro* (Bury and Sabo, 2011) and *in vivo* (Wu et al., 2013), suggesting that AZs might indirectly use KIF1A/UNC-104. Yet, more insights need to be gained on the mechanisms of AZ protein transport and recruitment to synapses.

To investigate the processes of both SV and AZ transport during synaptogenesis, several studies have taken a live imaging approach to track the recruitment of SVs (Ahmari et al., 2000), and AZs (Friedman et al., 2000) to the developing synapse in hippocampal cell cultures. These studies found that synapses form and become capable of synaptic vesicle release very quickly, in tens of minutes. Similarly, using an *in vivo* approach to observing synaptogenesis in Zebrafish, Jontes et al found that synaptic varicosities emerged in less than one hour at contact points between Mauthner cell axons and target post-synaptic cells (Jontes et al., 2000). Subsequently, several studies used similar *in vivo* approaches to examine the order of sequential recruitment of SVs, PTVs, and other synaptic components to the synapse (Böhme et al., 2016; Dai et al., 2006; Easley-Neal et al., 2013; Patel et al., 2006). In most cellular contexts, Liprin-□SYD-2 arrives at nascent synapses early in development (Dai et al., 2006; Fouquet et al., 2009; Patel et al., 2006), with Bruchpilot/ELKS-1 and RBP (Böhme et al., 2016) arriving later in a subsequent maturation step and being required for subsequent clustering of Calcium channels and pre-synapse maturation.

Despite this progress, the mechanisms underlying *en passant* synapse formation behind an extending growth cone *in vivo* are still not well understood. Nor is it known how dopamine synapses develop, and what characterizes their synaptic terminals. In this work, we developed an *in vivo* system to observe the *de novo en passant* synapse formation process using the *c. elegans* PDE dopamine neuron. Using this system, we find that dopaminergic PDE neurons develop bona fide Active Zones (AZs) at their synapses. We also find that AZ proteins and SVs coalesce into nascent synaptic puncta extremely rapidly behind the extending PDE growth cone within several minutes. Following this initial phase of rapid puncta formation, there is a second phase of synapse remodeling and organization where SVs and AZs increase in size and in puncta stability. We find that AZ proteins are mostly independent of UNC-104/Kinesin-3 for transport to the synapse; rather, our data suggest their transport by *unc-104*-independent molecular motors and/or diffusion. Finally, we show that *de novo* synapse formation depends on a proper balance between motor-driven transport and AZ dependent aggregation.

## Results

### Volume-transmitting dopamine neurons have bona fide Active Zones

PDE neurons are two bilaterally symmetric dopaminergic neurons with their cell bodies in the posterior mid-body of the worm. PDE neurons sense bacteria through their ciliated sensory endings that extend into the cuticle, and along with ADE and CEP dopamine neurons mediate the basal slowing response, a behavior which describes the worm slowing its locomotion when contacting bacteria (Sawin et al., 2000). More recently, dopamine has been shown to be necessary for precisely adjusting the locomotion velocity of worms, with mutants having greatly exaggerated velocity changes; again, exogenous application of dopamine was able to rescue this locomotion defect in worms (Omura et al., 2012).

The fact that one can rescue the basal slowing response (Sawin et al., 2000) and the ability to fine-tune velocity by bathing the worms in dopamine (Omura et al., 2012) suggests that precise neurotransmission at well-defined synaptic junctions is not necessary for behavioral rescue, which is consistent with the idea of a volume-transmission mode of dopamine release. PDE’s main downstream synaptic partner is the DVA interneuron, which expresses the D1-like dopamine receptor DOP-1C, and mediates PDE’s effects on the basal slowing response through downstream motor neurons (Bhattacharya et al., 2014). We therefore asked whether the anatomy of the synaptic connection between PDE and its downstream neuron, DVA, is supportive of volume transmission. We expressed a tdTomato-tagged DOP-1C receptor in DVA interneurons concurrently with a SNB-1::YFP transgene in PDE dopamine neurons, and observed that SNB-1::YFP forms discrete puncta along the PDE axon, while tdTomato::DOP-1C is diffuse along the DVA neurite (Figure 1B, C). The lack of precise alignment between pre and postsynaptic specializations at PDE synapses is reminiscent of dopaminergic synapses in vertebrates, where dopaminergic synaptic specializations have been observed at sites along the axon that are not in close apposition to a post-synaptic density and spine (Pereira et al., 2016; Rice and Cragg, 2008; Trudeau et al., 2014; Zhang et al., 2015).

**Figure 1:**
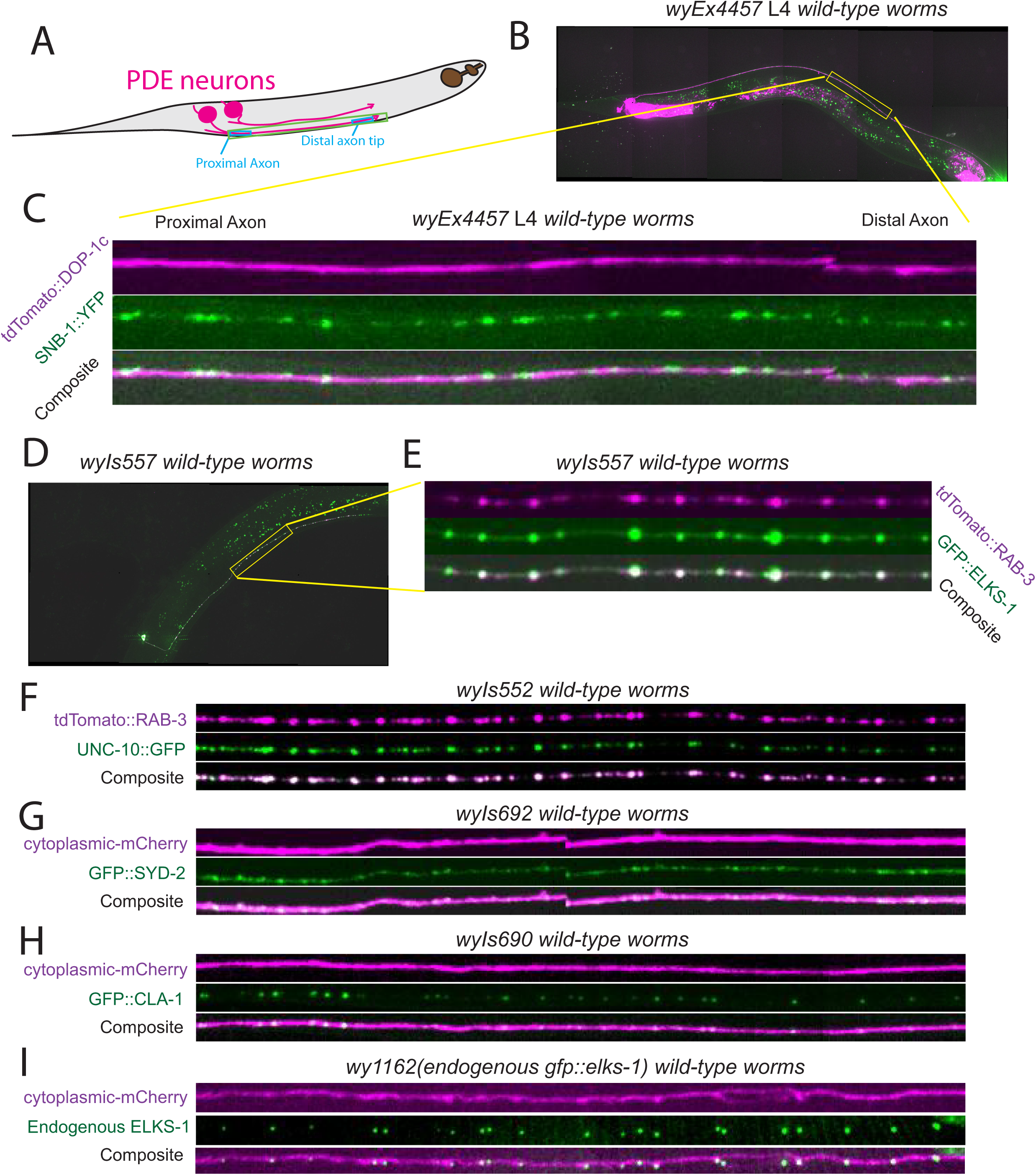
Volume-transmitting dopamine neurons have bona fide Active Zones. A) Model of two PDE neuronal cell bodies (shown in magenta) and their axons extending anteriorly along ventral nerve cord of a worm. B) Micrograph of *wyEx4457* L4 *wild-type* worm showing expression of tdTomato:DOP-1c using DVA-specific promoter and Pdat-1 driven SNB-1::GFP. C) Inset in Panel B. Shown is overlap of tdTomato::DOP-1c in DVA neurite, and SNB-1::YFP in PDE axon. D-E) Micrograph of *wyIs557* L4 *wild-type* worms, showing Pdat-1::tdTomato::RAB-3 and Pdat-1::GFP::ELKS-1, both expressed in PDE neurons. E) Inset in Panel D. F-H) Axon insets similar to Panels C and E depicting the following protein-XFP fusion constructs, which were expressed in PDE neurons using the Pdat-1 promoter: F) tdTomato::RAB-3 and UNC-10::GFP. G) cytoplasmic::mCherry and GFP::SYD-2. H) cytoplasmic::mCherry and GFP::Cla-1. I) Axon linescan from a worm expressing endogenously tagged GFP::ELKS-1 exclusively in dopamine neurons using Pdat-1 promoter in *wy1162*; *wyIs834*;+ Pdat-1::mCherry worms. *wy1162* allele = *elks-1 5’ utr::frt::gfp::pest-degron::sl2::frt::elks-1 5’ cds*. *wyIs834* = *Pdat-1::flippase*. Genomic locus modified using SapTrap kit (Schwartz et al. 2016).

Given this similarity between the PDE-DVA dopamine synapse and the vertebrate dopamine synapses, we wanted to ask whether such dopamine synapses contain bona fide active zones, as there is scant evidence of the molecular composition of dopaminergic pre-synaptic terminals (Pereira et al., 2016; Rice and Cragg, 2008; Trudeau et al., 2014; Zhang et al., 2015). We asked how key Active Zone molecules localize in PDE neurons, whether or not they formed discrete puncta in these neurons, and the extent to which they clustered with SV markers, focusing on the AZ proteins, SYD-2/Liprin-alpha, UNC-10/RIM and ELKS-1. SYD-2/Liprin-α is thought to be a key organizer of synapse assembly, which binds other AZ proteins, and recruits both additional AZ proteins and SVs to the synapse (Dai et al., 2006; Kittelmann et al., 2013; Patel et al., 2006; Zhen and Jin, 1999). UNC-10/RIM is another important AZ protein, which has been shown to play a crucial role in binding and tethering vesicles and pre-synaptic calcium channels (Kaeser et al., 2011; Südhof, 2012). Finally, ELKS-1 is an important synaptic scaffolding molecule, which binds SYD-2 (Dai et al., 2006; Kittelmann et al., 2013) and UNC-10 (Ohtsuka et al., 2002; Wang et al., 2002) and helps recruit calcium channels to the synapse (Kittel et al., 2006). As a SV marker, we used RAB-3, which is an abundant, small GTPase present on synaptic vesicles (Takamori et al., 2006) that helps recruit vesicles to the synapse through interactions with RIM (Südhof, 2013).

First, we co-expressed GFP::ELKS-1 and tdTomato::RAB-3 in PDE neurons, and found that both protein markers exhibited clear punctate localization patterns, and that there was extensive co-localization between both the SV marker RAB-3, and the AZ protein, ELKS-1 (Figure 1D, E). Next, we examined the fluorescence pattern of UNC-10/RIM and RAB-3, and found that they formed puncta, which also yielded a high degree of co-localization (Figure 1F). Expressing SYD-2 in PDE, we found that SYD-2 is present in very small but discrete puncta in PDE neurons. Finally, we assayed whether the AZ component, CLA-1, was present at these active zones in PDE. CLA-1 is a novel AZ component in *c. elegans*, recently characterized in our lab (Kurshan et al. personal communication) that is homologous to FIFE in *Drosophila*, as well as to Piccolo in mammals. Indeed, we found that CLA-1 also forms puncta when expressed in PDE dopamine neurons (Figures 1H). To confirm that AZ proteins are endogenously expressed in PDE neurons, we N-terminally tagged endogenous ELKS-1 in a cell-type specific manner using a conditional strategy based on flippase mediated recombination (Schwartz and Jorgensen, 2016). Here, Pdat-1 driven expression of a Flippase gene specifically in dopamine neurons catalyzes a DNA-excision event that results in GFP-tagging of the endogenous ELKS-1 gene. As can be seen, GFP::ELKS-1 puncta are clearly visible along the mCherry-marked PDE axon (Figure 1I). Thus, we conclude that PDE dopamine neurons contain canonical active zones that are defined by typical presynaptic cytomatrix molecules.

### Synaptic Vesicles (SVs) and Active Zone proteins (AZs) arrive at synapse quickly behind growth cone

Having observed that PDE neurons contain bona fide active zones, we wanted to ask how these dopaminergic *en passant* pre-synaptic terminals develop. To characterize the formation of these presynaptic specializations *in vivo*, we examined the synapse formation process behind the extending growth cone in PDE (Figure 2A, B). Shortly after PDE’s neurogenesis, during the L2 larval stage, PDE extends its axon towards the ventral nerve cord (VNC). Upon reaching the VNC, the PDE axon bifurcates and its major branch turns anteriorly and grows within the VNC, elaborating approximately 40 *en passant* synapses. Tracking the development of PDE, we found that the outgrowth of the PDE axon in the VNC was mostly completed within approximately 10 hours.

**Figure 2:**
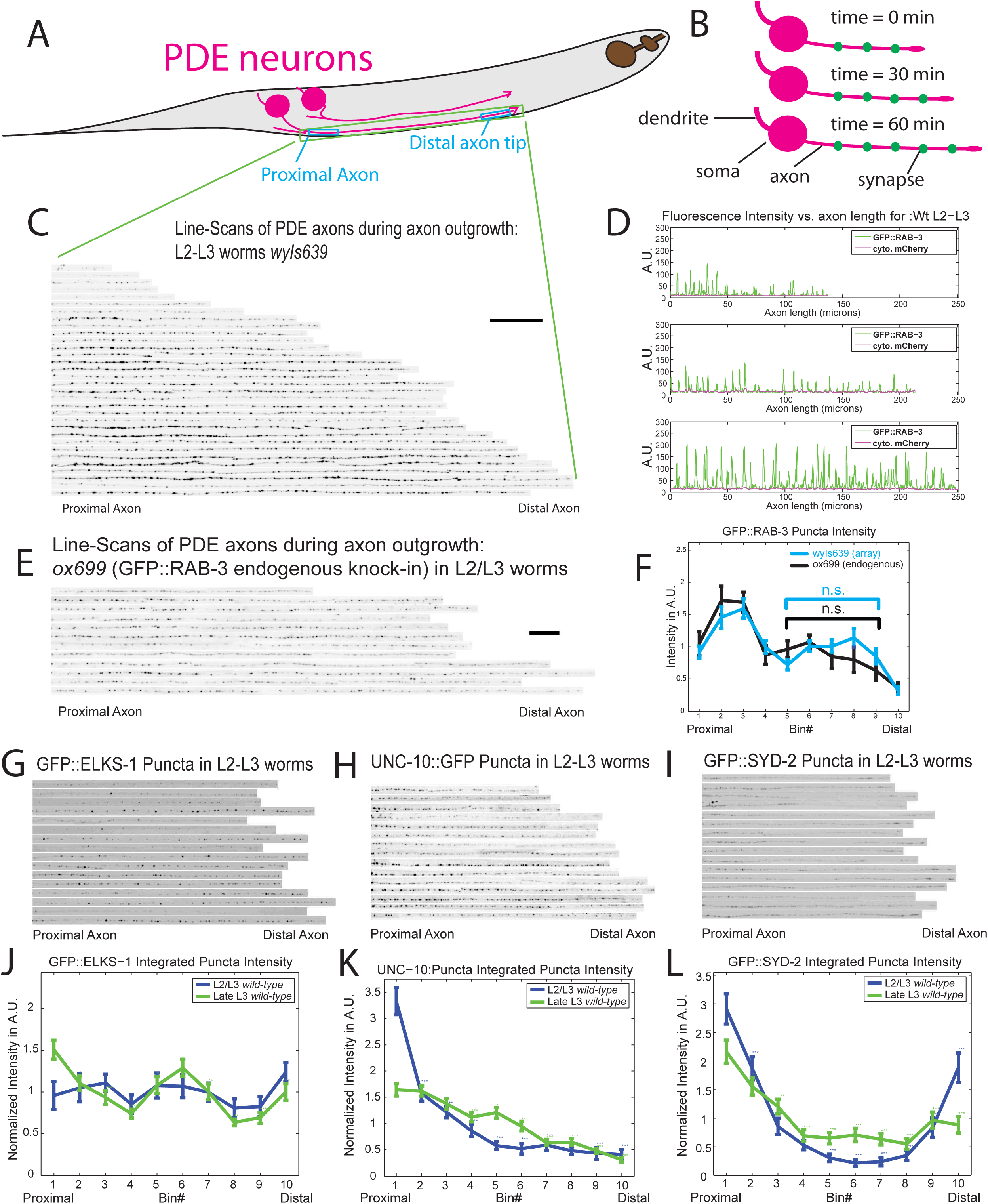
PDE neuron axonal outgrowth and synapse formation: an in vivo model of de novo synaptogenesis. A) Model of two PDE neuronal cell bodies (shown in magenta) and locations of proximal and distal axon segments. B) Model of PDE axon outgrowth over time. C) Line scans showing GFP::RAB-3 intensity in *wyIs639* wild-type worms as a function of PDE developmental stage, as measured by PDE axon length. Scale bar = 20 um. D) Plot profiles of GFP::RAB-3 and cytoplasmic-mCherry intensity along PDE axons. E) Line-scans of endogenously tagged GFP::RAB-3 in *ox699* worms with PDE-specific Flippase (Pdat-1::FLP). F) GFP::RAB-3 puncta distribution in *ox699* worms along PDE axons, proximal to distal. Axon was divided into 10 bins (1 is most proximal bin, and 10 is most distal) and thresholded to isolate puncta. Then, average summed puncta intensity was computed per bin. One way ANOVA test used to determine significance. Multiple comparisons Tukey’s h.s.d. test used to compare individual bins 2-10 for significant difference compared to bin 1. Significance values are computed only within a genotype cohort. Means and S.E.M.’s are shown. * = p < .05; ** = p < .01; *** = p < .001. G-I) Linescans showing puncta distribution along wild-type PDE axon in late-L2/early-L3 worms for the following Active Zone markers: G) GFP::ELKS-1 *(wyIs557)* H) UNC-10::GFP *(wyIs552)* I) GFP::SYD-2 *(wyIs692)*. J-L) Quantification of integrated puncta intensity in 10 binned segments, proximal to distal, of PDE axons. J) GFP::ELKS-1: n=38 L2/L3 axons and 34 late L3 axons. K) UNC-10::GFP: n=23 L2/L3 axons and 35 late L3 axons. L) GFP::SYD-2: n=34 L2/L3 axons and 39 late L3 axons. J-L) Statistical significance is result of comparisons between the first bin and all subsequent bins within an age cohort. One way ANOVA test used to determine significance. Multiple comparisons Tukey’s h.s.d. test used to compare individual bins 2-10 for significant difference compared to bin 1. Means and S.E.M.’s are shown. * = p < .05; ** = p < .01; *** = p < .001.

To determine how synapses form in these PDE axons, we co-expressed both a synaptic vesicle marker, GFP::RAB-3, and cytoplasmic mCherry under a dopamine neuron specific promoter, P*dat-1 (wyIs639)* and imaged these transgenic worms at different time points of axon outgrowth. Images of axons from many individual *wyIs639* worms were stacked together and aligned with the proximal axon on the left and the distal axon on the right (Figure 2C). Additionally, we randomly selected three of these axons and plotted the data as line scans of fluorescence intensity of GFP::RAB-3 and mCherry for each worm (Figure 2D). These analyses showed that we can reliably detect GFP::RAB-3 puncta within a few microns from the tip of the growing axon, suggesting that synaptic puncta form very rapidly behind the growth cone (Figure 2D). To quantify the distribution of SVs along the axon, we divided each PDE axon into 10 equal-length bins and computed the integrated puncta intensity of GFP::RAB-3 in each bin (Figure 2F). A relatively even distribution from bins 6-9 was observed, suggesting that the initial stage of synapse formation in PDE neurons is a rapid process. After this first phase of rapid synapse growth, a more gradual increase in synaptic vesicle intensity at mature synapses can be seen as the worm matures through L2/L3 to late L3 stage.

These data on synaptic vesicle distribution were obtained with a genome-integrated exogenously expressed transgenic array (*wyIs639*). To confirm that our observations reflected the endogenous RAB-3 distribution, we examined the endogenous RAB-3::GFP distribution using a cell-type specific endogenous GFP knock-in strain (*ox699*), in which the tagging of RAB-3 with GFP at the endogenous locus was activated by the expression of a *Pdat-1*::flippase (*wyEx8715*), as previously described (Figure 1I (Schwartz and Jorgensen, 2016)). We found that the endogenous RAB-3 distribution was very similar to that of the transgene (Figure 2E, 2F). Interestingly, the intensity of the endogenous GFP::RAB-3 was comparable to transgenic GFP::RAB-3 (data not shown), indicating that we were not overexpressing the RAB-3 marker.

Having observed the accumulation of SVs behind the growth cone, we asked how early during synaptogenesis AZs arrive at the synapse. To determine the time course of arrival of various AZ proteins, we imaged the distribution of ELKS-1::GFP (*wyIs557*), SYD-2/Liprin-α::GFP (*wyIs692*), and UNC-10/RIM::GFP (*wyIs552*) in PDE axons during axonal outgrowth and later developmental stages.

We took images of each of the following AZ proteins: ELKS-1, UNC-10 or SYD-2 fused to GFP in PDE axons. Images were taken during late L2 – mid L3 stages during which PDE extends its axon and forms synapses, and during late L3, when the axon outgrowth ends. We then formed axonal line scan images, similar to Figure 2C, for the AZ proteins ELKS-1, UNC-10, and SYD-2 (Figure 2G-I). All AZs examined were present along the axon at newly forming synaptic puncta from the very earliest time-points in PDE axon extension. However, upon examining the line-scans, there were some noticeable differences in the distribution of the AZs along the axon throughout development, suggesting that although all AZs were present at the earliest stages of synapse formation, they might accumulate at nascent synapses with different kinetics. For instance, SYD-2 is present in large amounts in the proximal axon and intriguingly has a distal tip accumulation as well (Figure 2I, 2L). This attenuates to a more even distribution as the worms proceed through development.

### Rapid accumulation of synaptic vesicle and active zone proteins behind the growth cone tip

To gain an even more detailed understanding of the rapid synapse formation process in the dopaminergic PDE neuron, we set out to observe in real-time and *in vivo* the *de novo* synapse formation events. We wanted to answer three questions: 1. Which type of pre-synaptic component arrives at these synapses first, vesicles or active zone proteins? 2. How quickly do both synaptic vesicles and active zone proteins accumulate and then coalesce at nascent synapses? 3. How variable are the arrival and coalescence events that lead to synapse formation? To measure synaptic vesicle accumulation, we used tdTomato::RAB-3, as mentioned previously. To track the accumulation of AZ components, we measured the intensity of GFP::ELKS-1.

Using time-lapse movies from transgenic worms exogenously expressing ELKS-1::GFP and TdTomato::RAB-3 (*wyIs557*) (Figure 3A), we generated kymographs to depict the intensities of both tdTomato::RAB-3 and GFP::ELKS-1 along the same axon over time (Figure 3B, C, D). Using the slope made by tracking the leading edge of the axon over time on the kymograph we were able to calculate the rate of axon outgrowth, which we observed to occur at rates as fast as 15 um/hour. This is consistent with the expected rate of axonal outgrowth given the distance the PDE axon must travel during the developmental time-window in which it extends (∼10 hours from mid-late L2 larval stage to mid-late L3 larval stage at 20° C). Using the kymographs we first asked how rapidly puncta were forming behind the extending axon. We find that GFP::ELKS-1 and tdTomato::RAB-3 cluster very quickly behind the extending growth cone (Figure 3B, C, D), in most cases forming synapses comparable in intensity to the mature synapses found along the axon base within 5-15 minutes. In addition, both ELKS-1 and RAB-3 cluster at these nascent synapses behind the growth cone at nearly the same time; neither protein arrived noticeably before the other one did. Our findings are not just limited to ELKS-1 as we also find that UNC-10/RIM and RAB-3 behave similarly, coalescing and aggregating together rapidly behind the growth cone (Supplementary Figure 3D).

**Figure 3:**
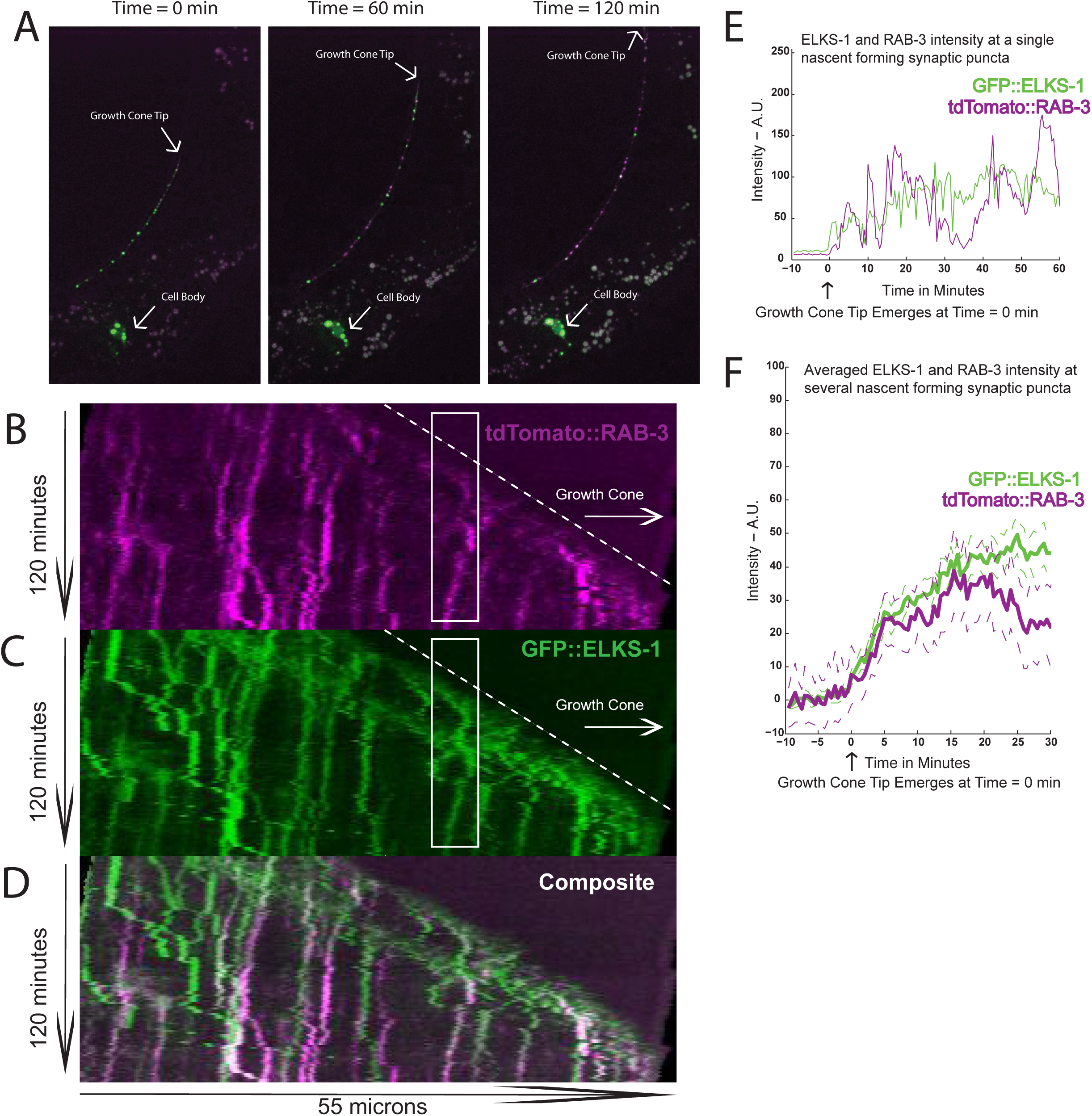
ELKS-1::GFP and tdTomato::Rab-3 coalesce quickly behind out-growing PDE axon tip. A) Images showing the PDE cell body and axon in *wyIs557 wild-type* worms at selected time-points during axon outgrowth. GFP::ELKS-1 depicted in green, tdTomato::RAB-3 in magenta. B-D) Kymographs showing tdTomato::RAB-3 and GFP::ELKS-1 fluorescence intensity in *wyIs557 wild-type* worms along out-growing PDE axon. Kymograph dimensions: Time = 120 minutes. Total distance = 55um. B) tdTomato::RAB-3 intensity. C) GFP::ELKS-1 intensity shown for same kymograph as in panel A. D) Composite kymograph showing both tdTomato::RAB-3 and GFP::ELKS-1 intensity. E) Fluorescence intensity of GFP::ELKS-1 and tdTomato::RAB-3 at a single developing puncta over time. An example puncta is highlighted by white box in panels B-D. ROI was selected by tracing backward in time from a stable GFP::ELKS-1 puncta. Both GFP::ELKS-1 and tdTomato::RAB-3 fluorescence intensities were then tracked at that segment of PDE axon, demarcated by ROI. F) Average of synapse formation intensity traces depicted in panel E. Dashed green and magenta plotted lines represent +/- S.E.M.’s of mean GFP::ELKS-1 and mean tdTomato::RAB-3.

To quantify the time course of synapse formation, and to assess variability in the process, we analyzed individual puncta formation events captured in the kymographs. An example of such a formation event can be seen in the white inset box within the kymographs (Figure 3B, C, D). A plot of ELKS-1 and RAB-3 fluorescence intensity reveals how quickly synapses can form (Figure 3E). For the synaptic punctum depicted in Figure 3 B-E, approximately 50 percent of its total intensity was gained in the first two minutes, with almost all of the rest of the increase occurring in the next 15-20 minutes. However, there was considerable variability in the rate of formation across various puncta formation events. Therefore, we averaged the fluorescence intensities of all newly forming puncta, and saw that both ELKS-1 and RAB-3 coalesce most rapidly in the first 5-10 minutes after the growth cone passes (Figure 3F). Then, there is a subsequent build-up over the next 10-15 minutes, completing this initial phase of synapse assembly behind the growth cone.

From the kymographs (Figures 3B-D), it appears that many of the RAB-3 puncta and the ELKS-1 puncta colocalize from their earliest appearance, indicative of ELKS-1 and SV aggregation, although the extent of colocalization does appear to increase with both time and distance from the leading edge of the growth cone. Indeed, thresholded images of both the ELKS-1 and RAB-3 channels of the long-term kymographs show considerable co-localization at the earliest time-points, with a subsequent increase as the puncta mature (Supplemental Figure 3A). We also quantified the colocalization of ELKS-1 and RAB-3 puncta throughout development using thresholded steady state images of *wyIs557* axons during development, again finding an initial phase of rapid colocalization followed by a second phase of maturation throughout the L3 stage, in which puncta continue to coalesce (Supplemental Figure 3B).

### Balance between transport and aggregation is crucial for correct positioning of synaptic vesicles, but not active zone proteins during initial synapse development

Having observed the rapid development of synapses in real-time in PDE, we asked what cell biological and molecular mechanisms determine how these dopaminergic synapses form. Previously we found that disrupting motor-driven transport in the cholinergic DA9 neuron by mutating either *unc-104/kinesin-3,* or its upstream activator, *arl-8,* disrupted SV distribution along the axon (Klassen et al., 2010; Niwa et al., 2016; Wu et al., 2013). Loss of the AZ-assembly molecule *syd-2* resulted in a similar phenotype as *unc-104/kinesin-3* gain-of-function mutants: reduction of SV clustering at synapses and the dispersal of SVs throughout axon. Each of these mutants could suppress the large ectopic SV clusters in the proximal axon in *arl-8(wy271)* mutants (Klassen et al., 2010; Wu et al., 2013). Taken together, these data suggest a model where motor-driven transport works antagonistically with synapse assembly-promoting active zone proteins to control vesicle aggregation and synapse formation (Klassen et al., 2010; Maeder et al., 2014; Wu et al., 2013).

However, these data are all from neurons with a classical asymmetric chemical synapse, consisting of a pre-synaptic bouton opposing a post-synaptic density. We wanted to ask how synapses are patterned in the volume transmitting dopaminergic PDE neuron. In addition, these previous data were obtained from mature axons, well after the synaptogenesis phase has been completed. It could be that synapses are initially laid down independently of motor-driven transport, and that subsequent growth and maturation processes require the proper coordination of motor-driven transport - as is the case for ELKS-1 homologue, Bruchpilot accumulating at *Drosophila* NMJ synapses (Pack-Chung et al., 2007). Therefore, we wanted to know how motor-driven transport mechanisms might affect the accumulation of both SVs and AZs at developing *en passant* dopamine synapses in the PDE neuron, at the exact time that they are first forming.

We first asked how reducing the strength of motor-driven transport by loss of *unc-104/kinesin-3* activator, *arl-8*, would affect the deposition of synaptic vesicles at developing synapses. In *wyIs557 wild-type* PDE neurons in L2/L3 worms, both the SV marker RAB-3 and the AZ protein ELKS-1 are evenly distributed along the axon. Loss of *arl-8* results in SV accumulation in the most proximal regions of the axon, concomitant with a loss of synaptic materials from the middle and distal axonal regions (Figure 4C, 4I). However, the distribution of ELKS-1 is unchanged in *arl-8* mutants (Figure 4D, 4J), suggesting that while SVs are dependent on *arl-8* mediated activation to reach the synapse, AZs such as ELKS-1 do not require ARL-8 mediated UNC-104/Kinesin-3 transport.

**Figure 4:**
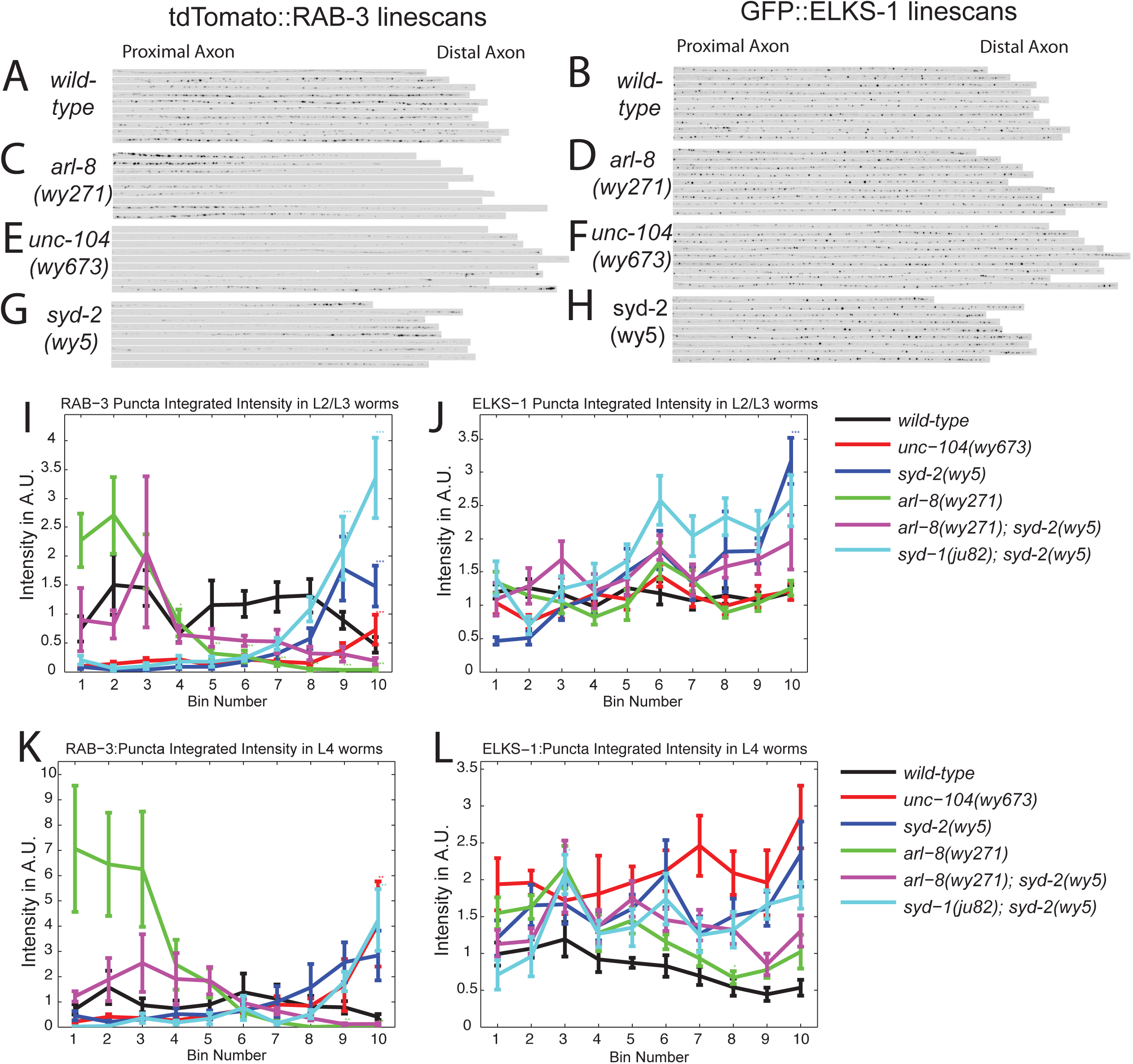
Balance between transport and aggregation is crucial for correct positioning of synaptic vesicles, but not active zone proteins during initial synapse development. A-J) Linescans of wyIs557 L2-L3 worms for the following genotypes wild-type, unc-104(673), syd-2(wy5), arl-8 (wy271). A, C, E, G) tdTomato::RAB-3 in *wyIs557* L2-L3 worms. B, D, F, H) GFP::ELKS-1 in the same *wyIs557* L2-L3 worms as A, C, E, G. I-L) Integrated puncta intensity per bin line plots in wyIs557 worms for the following genotypes: *wild-type*, *unc-104(673)*, *syd-2(wy5)*, *arl-8 (wy271)*, *arl-8 (wy271); syd-2(wy5), syd-1(ju82); syd-2(wy5)*. I-J) tdTomato::RAB-3 and GFP::ELKS-1 intensity plots in L2/L3 worms. Number of worms: wild-type n = 23, *unc-104(673)* n = 25, *syd-2(wy5)* n = 15, *arl-8(wy271)* n = 20, *arl-8 (wy271); syd-2(wy5)* n = 7, *syd-1(ju82); syd-2(wy5)* = 17. K-L) tdTomato::RAB-3 and GFP::ELKS-1 intensity plots in L4 worms. Number of worms: *wild-type* n = 17, *unc-104(673)* n = 9*, syd-2(wy5)* n = 9, *arl-8(wy271)* n = 10, *arl-8(wy271); syd-2(wy5)* n = 12, *syd-1(ju82); syd-2(wy5)* n = 5. I-L) One way ANOVA test used to determine significance. Subsequently, multiple comparisons Tukey’s h.s.d. test used to compare individual bins 2-10 for significant difference compared to bin 1. Significance values are computed only within a genotype cohort. Means and S.E.M.’s are shown. * = p < .05; ** = p < .01; *** = p < .001.

In the *unc-104(wy673)* mutant, where auto-inhibition is prevented, leading to over-activation of Kinesin-3 (Niwa et al., 2016) SVs are dramatically reduced at proximal and medial regions along the axon but remain present near the tip of distal axon (Figure 4E, 4I). This severe phenotype argues that the auto-inhibition of UNC-104 is a regulatory mechanism that is essential for PDE synapse formation. Interestingly, in this overactive Kinesin-3 mutant, the ELKS-1 distribution pattern is still unchanged (Figure 4F, 4J), arguing that ELKS-1 proteins do not depend on regulation of Kinesin-3 activity for transport to synapses.

Next, we asked how synaptic assembly promoting AZ proteins might recruit transiting SVs and AZ components to nascent synapses. Loss-of-function of the active zone protein *syd-2 (wy5)* resulted in a reduction of SV intensity at synapses along most of the axon, except for the most distal segments, where they accumulated (Figure 4G, 4I). *Syd-1 syd-2* double mutants exhibited a similar phenotype to the *syd-2* single mutant (Figure 4H, 4J). In *syd-2* single and *syd-1 syd-2* double mutants, ELKS-1 intensity slightly increased in the distal axon in L2/L3 worms (Figure 4H, 4J). This increase is transient, however; in L4 stage worms, the distal synapses contain similar amounts of ELKS-1 as do *wild-type* controls (Figure 4L). Thus, whereas manipulations of both motor-driven transport and localization of synapse-assembly AZ molecules affect SV distribution, ELKS-1 localizes to synapses mostly independently of both processes.

We then asked whether we could rescue defects in one of the two forces that affect SV localization (kinesin motor-strength or pro-assembly AZ molecules) by modulating the strength of the other pathway. Thus we made *arl-8 syd-2* double mutants, and found that the distribution of RAB-3 labeled SVs in these worms is shifted more distally as compared to the *arl-8 single* mutant alone. Again, ELKS-1 localization was unaffected (4E). These results suggest that motor-driven transport and pro-assembly AZ molecules have the same net output: to determine the deposition of SVs along the axon. Deposition of SVs at synapses both during development and during subsequent synapse growth thus likely requires three separate steps. 1. Activation of *arl-8* and transport of SVs out of the cell body and down the axon. 2. Deactivating kinesin-3 transport to halt transiting SVs at *en passant* pause sites along the axon. 3. Recruitment of SVs and incorporation into nascent synaptic puncta by pro-assembly molecules like *syd-2*.

Importantly, we observed almost identical phenotypes between L2/L3 worms and L4 worms for both SV distribution (Figure 4I, 4K) and for the distribution of the AZ protein, ELKS-1 (Figure 4J, 4L). This suggests that the same transport and AZ-dependent synaptogenic mechanisms operate both during *de novo* synapse formation during development, and during later synapse growth and maturation. These data also demonstrate that the same intra-cellular transport and active zone assembly mechanisms, which pattern cholinergic pre-synapses (Klassen et al., 2010; Wu et al., 2013), also function to instruct the development of volume-transmitting dopamine pre-synaptic terminals.

### Vesicle transport dynamically regulated by *unc-104*/kinesin-3 activation state and AZ proteins to deposit vesicles at synapses

To gain more insight into how active zone proteins and motor-driven transport coordinate the distribution of synaptic vesicle cargos in a developmental context, we measured the dynamic movement behavior of SVs in PDE in GFP::RAB-3 (*wyIs639) wild-type* and mutant animals in L3 stage worms. Here, we photobleached a region in the proximal PDE axon to reduce background fluorescence, and then took time-lapse movies of GFP::RAB-3, allowing us to visualize vesicular transport events, as well as the pausing frequency and the resident pause time of these SV transport events, all throughout the axon.

First, we found that *arl-8* mutants had dramatically reduced transport event frequency (Figure 5C, E), and reduced anterograde velocities (Figure 5G). There was also an increased pause time of transport events (Figure 5K). These data argue strongly that ARL-8 is required to activate UNC-104/Kinesin-3 mediated axonal transport during the synaptogenesis phase. These dynamic phenotypes satisfyingly explain the lack of synapses in the distal axon in the *arl-8* mutants. Consistent with this idea, over-activating Kinesin-3 motor transport efficacy with a gain of function mutation in *unc-104 (wy673)* resulted in an increase in run length of anterograde transport, suggestive of a defect in pausing frequency (Figure 5I). In addition, *unc-104 (wy673)* also showed a decrease of retrograde event numbers (Figure 5F), indicative of a disrupted balance between anterograde and retrograde transport.

**Figure 5:**
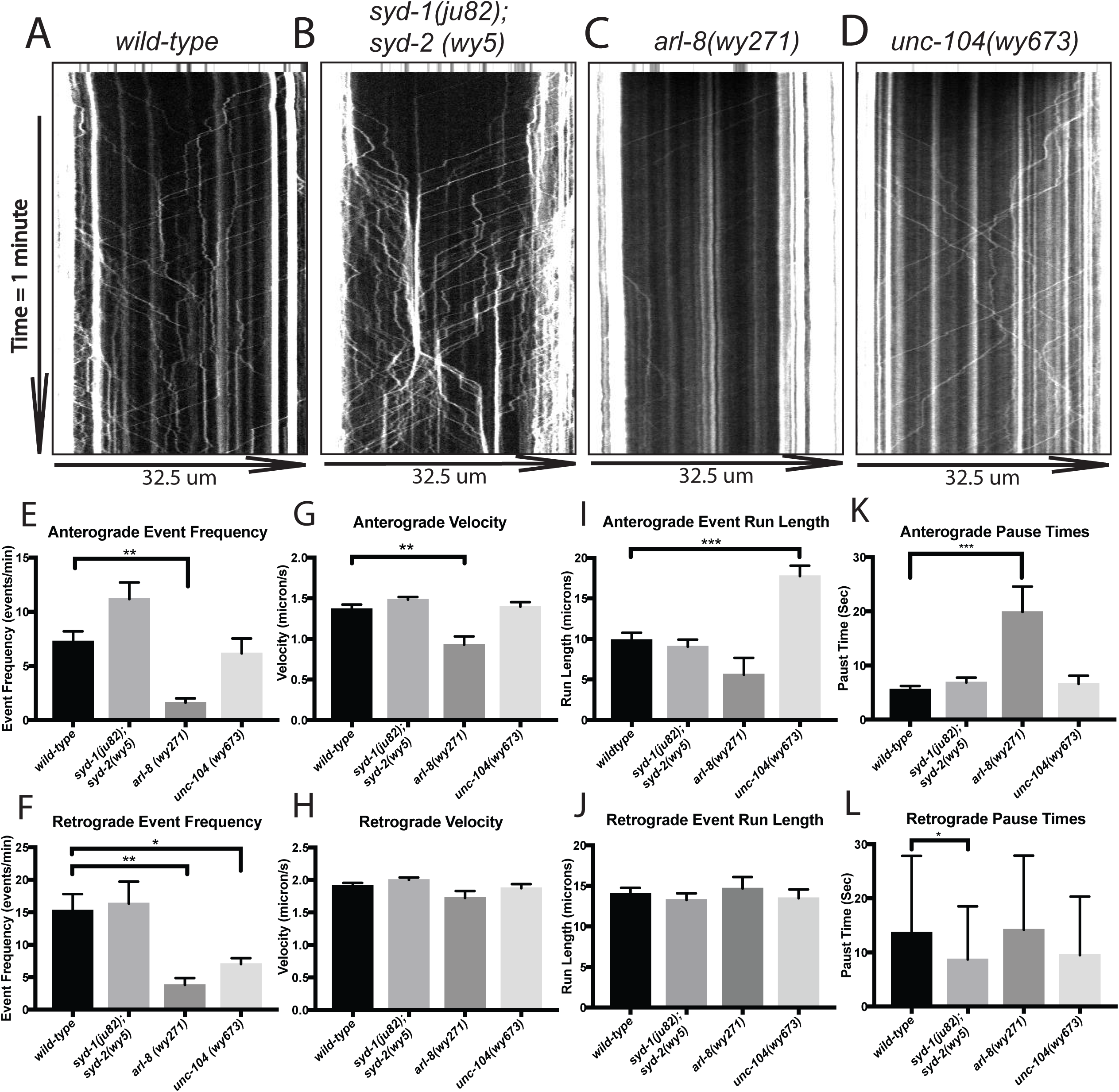
Vesicle transport dynamically regulated by unc-104/kinesin-3 activation state and AZ proteins to deposit vesicles at synapses. A-D) Kymographs showing GFP::RAB-3 transport in PDE axon base in *wyIs639* L3 worms (depicted in Figure 1A). Kymographs were taken over the course of 1 minute. Anterograde events are those traveling to the right, and retrograde to the left. Top 10 rows of kymograph represent the pre-bleach image of the axon. Kymographs taken after bleaching. A) *wild-type* B) *syd-1(ju82), syd-2(wy5)*. C) *arl-8 (wy271)*. D) *unc-104 (wy673)*. E-F) Retrograde and anterograde transport event frequencies, measured as the mean number of events observed per minute. *wild-type* n = 8, *syd-1(ju82); syd-2(wy5)* n = 6, *arl-8 (wy271)* n = 5, *unc-104 (wy673)* = 8. G-H) Anterograde and Retrograde velocities in um/sec. I, J) Anterograde and Retrograde run lengths, in um traveled from entering bleached imaging window until pausing. G, I) Anterograde event #’s: *wild-type* n = 59, *syd-1(ju82); syd-2(wy5)* n = 67, *arl-8 (wy271)* n = 7, *unc-104 (wy673)* = 43. H, J) Retrograde event #’s: *wild-type* n = 122, *syd-1(ju82); syd-2(wy5)* n = 98, *arl-8 (wy271)* n = 19, *unc-104 (wy673)* = 49. K) Anterograde pause time event #’s: *wild-type* n = 87, *syd-1(ju82); syd-2(wy5)* n = 100, *arl-8 (wy271)* n = 10, *unc-104 (wy673)* = 39. L) Retrograde pause time #’s: *wild-type* n = 112, *syd-1(ju82); syd-2(wy5)* n = 117, *arl-8 (wy271)* n = 16, *unc-104 (wy673)* = 52. Panels E-L) Significance determined by one way ANOVA (Fig E-H) or Kruskal-Wallis (I-L) tests followed by Dunnet’s multiple comparison’s test to compare mutant genotypes to wild-type. * = p < .05; ** = p < .01; *** = p < .001.

In *syd-1 syd-2* double mutants, mean run length and velocity were unaffected. However, we observed an increase in anterograde event frequency (Figure 5E) that was just short of significance. We have previously observed increased transport event frequencies in *syd-2* (*wy5*) mutants in another neuron, DA9 (Wu et al., 2013), suggesting that this event frequency increase in PDE may be real. In *syd-1 syd-2* double mutants we also observed a slight decrease in retrograde event pause time (Figure 5L). The fact that these AZ protein mutants did not affect velocity or run length suggests that the synapse-assembly promoting AZ proteins do not directly affect the activation state of the kinesin motor and/or motor processivity. Rather, the specific role that assembly-promoting AZ proteins might play is to help control the recruitment of paused transport packets to synaptic sites along the axon. Importantly, the effects of UNC-104, ARL-8, SYD-1 and SYD-2 on SV distribution during the synaptogenesis phase demonstrate that both regulation of SV transport and presynaptic assembly-inducing AZ molecules cooperate to pattern *en passant* synapses along the axon during synapse development.

### Active zone proteins and SVPs exhibit different dynamics of transport along axon

Having observed that SVs are highly dependent on *unc-104/kinesin-3* for transport along the axon we wanted to ask how AZs traffic to the synapse. Previously, we saw that manipulations, which affect Kinesin-3’s transport activity leave the distribution of the AZ protein, ELKS-1 largely unaffected (Figure 4), suggesting its lack of motor-dependent transport, at least by the Kinesin-3 motor. Yet, there is prior evidence that AZs travel along the axon by motor-driven transport (Klassen et al., 2010; Pack-Chung et al., 2007; Shapira et al., 2003), so we wanted to see if we could observe AZ transport behavior in PDE.

We thus took time-lapse movies of ELKS-1, UNC-10, or SYD-2 tagged with GFP to observe their dynamic behaviors, after first photobleaching the axon to reduce background intensity from stable puncta. Compared to RAB-3 (Figure 6D), all the AZ proteins had far fewer observable transport events, by at least an order of magnitude (Figure 6A-E). There were some slight differences in trafficking parameters between the AZ proteins. For instance, SYD-2 was notable for having transport events of very long run length (Supplemental Figure 6A). Overall, AZ proteins exhibit transport events that are very infrequent, especially compared to the numbers of SV events we observed. Based on their frequency, these observable transport events are unlikely to provide enough material for the formation of pre-synapses behind the axon growth cone tip, based on their frequency and intensity.

**Figure 6:**
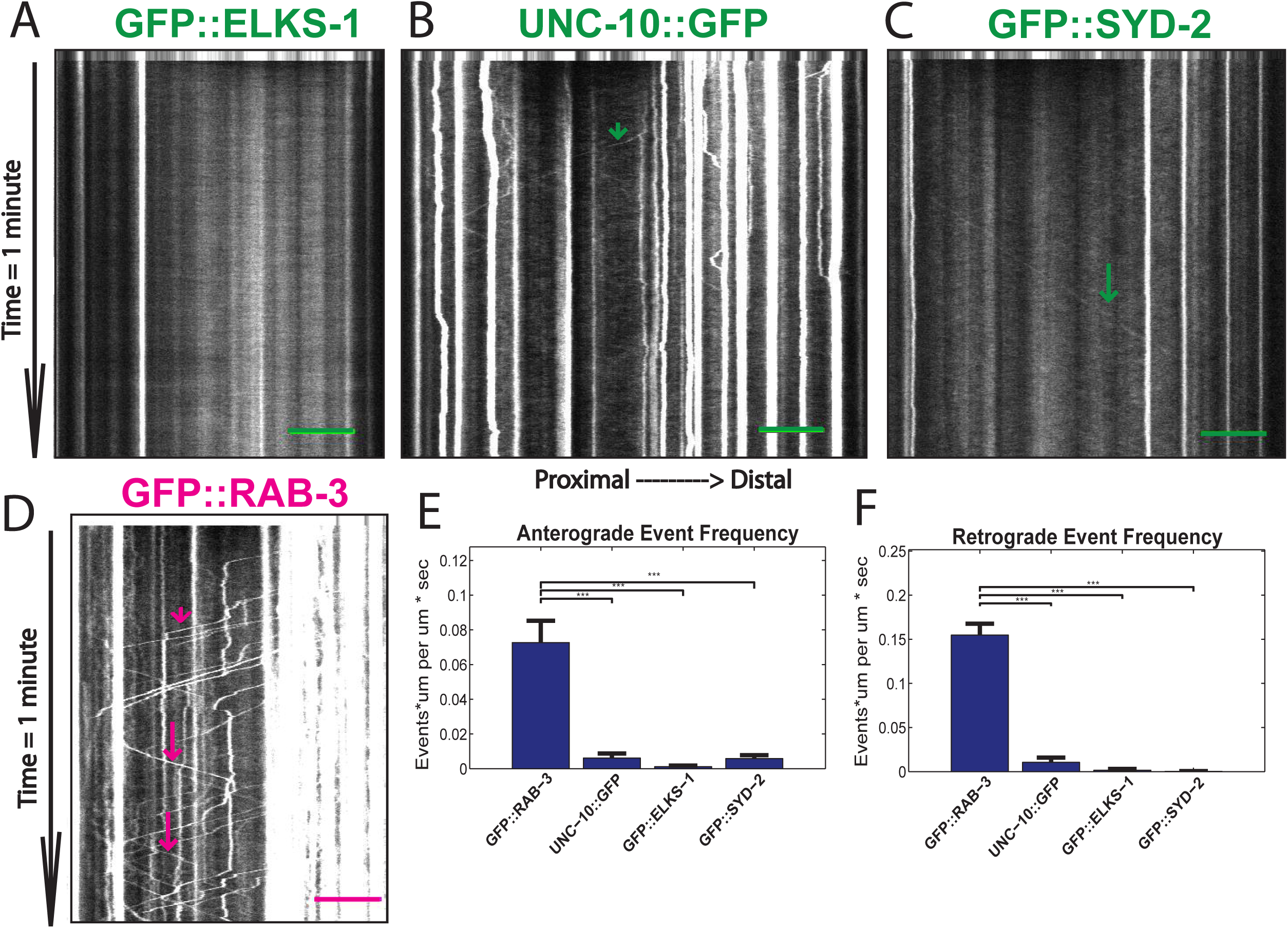
Active Zone proteins and SV’s exhibit different dynamics of transport along axon. A-D) Kymographs from streaming movies of protein localization along axon base in L3 worms. A) GFP::ELKS-1 (*wyIs557*). B) UNC-10::GFP (*wyIs552*) C) GFP::SYD-2 (*wyIs692*). D) GFP::RAB-3 (*wyIs552*). Scale bars are 5um. Kymographs are 1min long, with images taken at 7.5 Hz, or every 133 ms. Long arrows indicate anterograde events. Short arrows indicate retrograde events. The first 10 rows of the kymograph represent the pre-bleach image. E) Anterograde event frequency. F) Retrograde event frequency. E-F) Frequency values were calculated by summing total run lengths of all events imaged and dividing this sum by (total distance imaged multiplied by total time of imaging). N values are # of kymographs imaged: GFP::RAB-3: n = 17, UNC-10::GFP: n = 34, GFP::ELKS-1: n = 23, GFP::SYD-2: n = 29. E-F) Analysis of transport in kymographs of streaming movies of transport in proximal axon in L3-L4 stage worms. Kruskal-wallis test followed by multiple comparisons Tukey’s h.s.d. test used to compute significance of differences between each of individual bins 2-10 and bin 1. Means and S.E.M.’s are shown. * = p < .05; ** = p < .01; *** = p < .001.

### Active zone proteins have different dependencies on UNC-104/KIF1A for transport to synapses

Having observed different trafficking dynamics of active zone proteins as compared to synaptic vesicles, we asked whether AZs and SVs used different molecular mechanisms for axonal trafficking. It is known that synaptic vesicle precursors are dependent on UNC-104/Kinesin-3 for their transport to synapses (Hall and Hedgecock, 1991; Okada et al., 1995). However, it is not known how AZ proteins are transported to the synapse. Manipulations of microtubule-motor driven vesicle trafficking variably affects the distribution of active zone proteins. Even the same AZ protein is variably dependent on UNC-104/Kinesin-3 in different cell types (Goldstein et al., 2008; Klassen et al., 2010; Pack-Chung et al., 2007; Patel et al., 2006). Furthermore, the dependence on Kinesin-3 for synaptic localization is in some cases developmentally regulated (Pack-Chung et al., 2007). Thus, we wanted to ask how active zone formation is affected by loss of *unc-104/kinesin-3* in PDE neurons when synaptic active zones are first forming.

Whereas synaptic vesicle distribution is greatly perturbed by loss of function of *unc-104/kinesin-3*, ELKS-1 distribution seems to be largely unaffected in PDE in *unc-104/kinesin-3* mutants. Indeed, although RAB-3 clusters are still present in the most proximal part of the axon, we observed that RAB-3 was absent from most of the axon in *unc-104/kinesin-3* mutants (Figure 7A). In contrast, ELKS-1 distribution was uniform from proximal to distal segments of the axon in both *wild-type* worms and in *unc-104/kinesin-3* mutants (Figure 7A, C). Syd-2 was also *unc-104* independent: in *wild-type* worms, SYD-2 is naturally distributed in a proximal to distal gradient; in *unc-104/kinesin-3 (e1265)* worms, this distribution is largely unaffected (Figure 7D). In contrast, UNC-10/RIM showed a much greater dependence on UNC-104/Kinesin-3. In wild-type worms, UNC-10 shows a mild proximal-distal gradient. This gradient was visibly exacerbated in *unc-104/kif1a* mutant worms, mostly due to a marked increase in the amount of material in the most proximal part of the axon (Figure 7E). However, the UNC-10/RIM phenotype was not as severe as what is observed for the synaptic vesicle marker RAB-3 (Figure 7B), as distal regions still clearly had small UNC-10/RIM puncta. These data indicate that different active zone proteins may be differentially dependent on UNC-104 for transport: ELKS-1 and SYD-2 were largely independent of *unc-104/kinesin-3*, whereas UNC-10/RIM was much more dependent on this Kinesin-3 motor.

**Figure 7:**
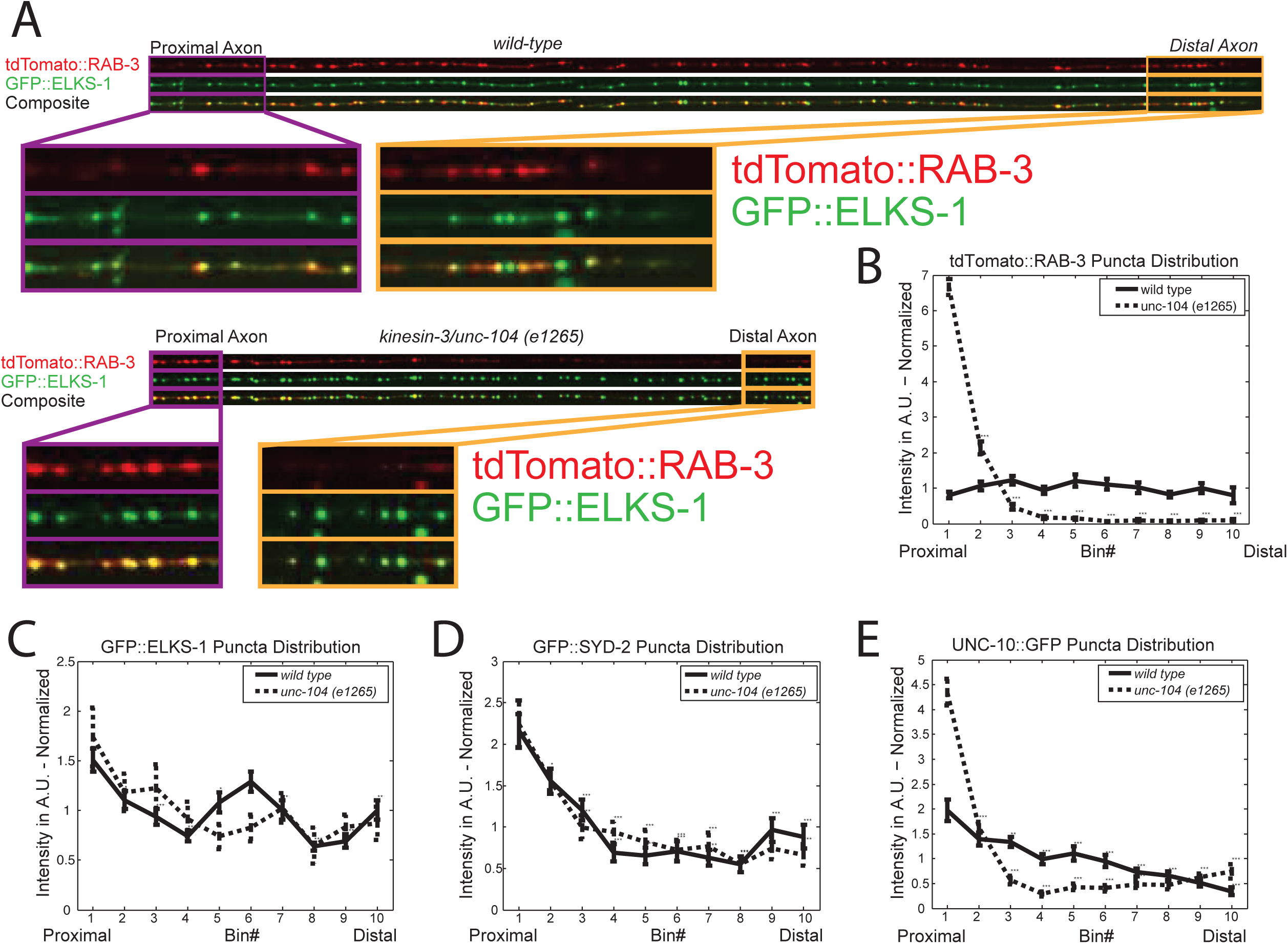
Active Zone proteins have different dependencies on Unc-104/Kinesin-3 for transport to synapses. A) Linescans of tdTomato::RAB-3 and GFP::ELKS-1 in wild type and *unc-104* (*e1265*) mutant worms. B-E) Distribution of B) RAB-3, C) ELKS-1, D) SYD-2 and E) UNC-10 puncta along PDE axon. Axons were divided into 10 bins (bin 1 most proximal, bin 10 most distal) and average puncta intensity was computed for each bin in both wild type and *unc-104* (*e1265*) mutants. N numbers are # of axons: B) tdTomato::RAB-3 in *wyIs552* worms. *wild-type*, n = 23, *unc-104* (*e1265*) n = 25. C) GFP::ELKS-1 in *wyIs557* worms. *wild-type*, n = 34, *unc-104 (e1265)* n = 12. D) GFP::SYD-2 in *wyIs692* worms. *wild-type*, n = 38, *unc-104 (e1265)* n = 12. E) UNC-10::GFP in *wyIs552* worms. *wild-type*, n = 23, *unc-104 (e1265)* n = 25. B-E) Statistical significance is result of comparisons between the first bin and all subsequent bins within a single genotype cohort (either *wild-type* or *unc-104 (e1265)*). One way ANOVA test used to determine significance. Subsequently, multiple comparisons Tukey’s h.s.d. test used to compare individual bins 2-10 for significant difference compared to bin 1. Significance values are computed only within a genotype cohort. Means and S.E.M.’s are shown. * = p < .05; ** = p < .01; *** = p < .001.

Previously, we observed very rare movements of GFP::ELKS-1 in the long-term movies of axon outgrowth and synapse formation that were reminiscent of motor-driven transport (Figure 3). We wanted to see if these rare rapid AZ protein movements were *unc-104/kinesin-3* dependent. Thus, we took long-term movies of synapse formation in L2/L3 *wyIs557 unc-104/kinesin-3 (e1265)* mutant worms. Surprisingly GFP::ELKS-1 exhibited transport events in these *unc-104/kinesin-3 (e1265)* mutant worms (Supplemental Figure 7), suggesting that they are not mediated by Kinesin-3.

## Discussion

Understanding how active zone components and synaptic vesicles assemble into presynaptic terminals is an important question in neuronal cell biology. While several important studies have examined this question in glutamatergic CNS or cholinergic NMJ synapses, all of which have a classic post-synaptic density opposing a pre-synaptic specialization, to our knowledge no study has examined the molecular mechanisms of synaptogenesis of volume-transmitting dopamine synapses. Also, while several important studies have tackled the question of synapse development *in vitro*, or *in vivo*, few have dissected the molecular mechanisms of *en passant* synapse formation *in vivo*, while observing the process of *de novo* synapse formation in real-time during development.

Tracking this developmental process *in vivo*, we find that the initial stages of synapse formation, characterized by the rapid and convergent aggregation of synaptic vesicles and active zone proteins behind the developing growth cone, occur in as little as a few minutes (Figures 2 and 3). After the initial synaptogenesis events, there is a subsequent phase of maturation and growth, characterized by a continued increase in the amount of certain AZ proteins and SVs present at the synapse, increased co-localization of AZ and SV markers, and increased stability of synaptic positioning (Figures 2 and 3).

While it has long been known that synaptic vesicle precursors require UNC-104/Kinesin-3 for transport to synapses (Hall and Hedgecock, 1991; Okada et al., 1995; Pack-Chung et al., 2007), it is not well known how vesicle transport is regulated to deposit SVs at nascent synapses. This is a conceptually interesting question, especially for *en passant* synapses: since there are many *en passant* synaptic sites along the axon that can potentially receive the transport cargoes, the problem of how to deposit SV cargos among these many sites is particularly acute in this context. Dopamine neurons, which are some of the most widely arborized neurons in the brain, having many thousands of *en passant* synapses (Matsuda et al., 2009), are especially challenged in this regard.

We find that two intracellular processes must occur to appropriately distribute transported SVs at synaptic sites along the axon. First, the UNC-104/Kinesin-3 motor is inhibited, allowing the transported cargos to pause when the Kinesin-3 motor has been switched to the off state. Second, pro-assembly AZ molecules are necessary to ensure that transiting vesicles are appropriately captured and retained at specific sites along the axon. Our evidence for the first part of this model is that over-activating the Kinesin-3 motor by making an intra-molecular mutation, which disrupts its auto-inhibition and results in the motor being preferentially in the ‘on’ state (Niwa et al., 2016), resulted in the over-accumulation of SVs at the axon tip and a dramatic decrease in synaptic vesicle accumulations at synapses along the length of the axon (Figure 4). Dynamic imaging revealed that transiting SVs paused less frequently, exhibiting a longer run length in these mutants (Figure 5). Similarly loss of Kinesin-3 activator, *arl-8*, which would leave the Kinesin motor preferentially in the ‘off’ state, resulted in pre-maturely aggregated puncta in the proximal axon, and little to no puncta distally. Dynamic imaging showed that transport events occurred much less frequently, and paused for a longer amount of time (Figure 5).

Meanwhile, consistent with the second part of our model, loss of presynaptic assembly molecules, including SYD-2 and SYD-1, causes a dramatic reduction of SV accumulation at synapses and an accumulation towards the distal axon tip (Figure 4), a phenotype that is somewhat similar to the overactive Kinesin-3 mutant. This phenotype is reminiscent of what has been observed in *Drosophila* motoneurons, where ectopic accumulations of vesicles are similarly found mis-localized in distal regions of the axon, proximal to the synaptic terminals (Li et al., 2014). In both cases, loss of *syd-2* and/or *syd-1* could result in transiting SVs bypassing their targets and accumulating at the nearest downstream location that motor-driven transport carries them. This would be consistent with a model of a conveyor belt of SV transport, where SVs are trafficked in multiple rounds along axons before accumulating at *en passant* synaptic sites, again similar to what has been observed for DCV trafficking in *Drosophila* motoneurons (Wong et al., 2012), as well as our findings from other neurons in *C. Elegans* (Patel et al., 2006; Wu et al., 2013).

In contrast to synaptic vesicles, how individual active zone proteins travel to the synapse is less well understood (Petzoldt et al., 2016). We therefore sought to shed some light on this process in PDE neurons. Despite the rapidity with which AZ molecules such as ELKS-1 accumulated at nascent synapses in PDE, very low frequencies of AZ protein movements were observed (Figure 6). These infrequently observed transport events are very likely insufficient to account for the rapid accumulation of AZ molecules at nascent synapses. It is possible that most PTV transport packets in PDE neurons contain very few copies of AZ proteins, and thus observation of the majority of transport events might require a more sensitive method of detection. Alternatively, it is possible that in PDE neurons some AZs aggregate at nascent synapses via passive diffusion. It is also possible that AZ proteins use a combination of both motor-driven transport and diffusion to travel to the synapse.

If AZ transport events do exist in PDE, and simply aren’t observable, then in addition to the possibility of their being transported on PTV’s, one further possibility is that AZ proteins hitch a ride on synaptic vesicles, as has been observed for synapsin-I in hippocampal neurons (Scott et al., 2011; Tang et al., 2013). Indeed, there has been co-trafficking observed between synaptic vesicles and active zone proteins in both cultured hippocampal neurons (Tao-Cheng, 2007), and in the c. elegans DA9 neuron (Wu et al., 2013). If this were the case that AZs were co-trafficked on SVs, then one would expect them to be dependent on Kinesin-3 for transport and localization to the synapse: Consistent with this idea, UNC-10 exhibited a moderate dependence on *unc-104/kinesin-3* for proper synapse distribution in PDE (Figure 7).

However, both SYD-2 and ELKS-1 localized to the synapse independently of UNC-104/Kinesin-3 in PDE neurons, and all the manipulations of Kinesin-3 efficacy, which dramatically altered SV localization patterns, left ELKS-1 puncta mostly indistinguishable from *wild-type* (Figure 4). In addition, in *unc-104* loss of function mutants, we still observe ELKS-1 transport-events in the long-term movies (Supplemental Figure 7A). These data suggest that ELKS-1 and perhaps other AZ proteins as well do not co-traffic with SVs down the axon.

Overall, our data on AZ localization and transport point toward the possibility that different AZ molecules use different mechanisms for transport to the synapse. The specific modes of transport for each of the AZ molecules might be determined by their position and function within the complex set of interactions that form the active zone. For instance, UNC-10/RIM, which binds more directly to synaptic vesicles through direct interactions with RAB-3 (Petzoldt et al., 2016; Südhof, 2012), exhibited some dependence on UNC-104/Kinesin-3 for localization to the synapse. In contrast, SYD-2 and ELKS-1, which bind SVs more indirectly, were independent of Kinesin-3/SVs for transport to the synapse in PDE. AZ proteins could also have dominant and secondary modes of transport. Indeed, vesicle-like movements were observed for SYD-2 (Figure 6) but in the absence of *unc-104*/*kinesin-3*, SYD-2 still localized to synapses perfectly well (Figure 7), suggesting that SYD-2 might be capable of both vesicular and non-vesicular transport.

Regardless of how AZ molecules travel down the axon, we observe that many of them naturally accumulate in proximal-distal gradients along the axon. This is true for UNC-10, SYD-2 (which also has a distal tip accumulation) (Figure 2), and Cla-1 (data not shown). Intriguingly, a recent study found that DCV’s are also distributed in a proximal-distal gradient in type III *Drosophila* motoneurons, which form a large number of *en passant* synapses along axonal branches (Tao et al., 2017). In this study, the DCV gradient attenuates over the course of development, as more materials are distributed distally. This is remarkably similar to the phenotype we observe in PDE neurons. Since it is currently hypothesized in the literature that AZs are transported on PTVs, a type of vesicle that looks very similar to DCVs by EM, this similarity in axonal distribution of AZs and DCVs is particularly interesting. SYD-2’s tip enrichment was an interesting and unexpected observation, and raises the intriguing possibility that there is a pool of SYD-2 molecules right behind the growth cone that are primed to initiate synaptogenesis. This possibility is especially interesting given that Syd-2 is thought to be an early initiator of synapse assembly (Dai et al., 2006; Patel et al., 2006).

In summary, we have established a model system for examining the process of *de novo* synaptogenesis of *en passant* volume-transmitting dopamine synapses along an axon *in vivo*. First, we characterize the molecular composition of PDE dopaminergic pre-synaptic terminals, and find that they are comprised of many canonical AZ proteins, which are well co-localized with SV puncta. We find that PDE synaptogenesis events occur rapidly, in as little as a few minutes. We also show that proper regulation of Kinesin-3 mediated axonal transport as well as the action of synapse-assembly promoting AZ molecules are necessary for appropriately depositing SVs at nascent *en passant* synapses during development. Finally, we show that while SVs are highly dependent on Kinesin-3 transport mechanisms to travel to the synapse, most AZ proteins travel along the cell body to the synapse in PDE neurons largely by other means.

## Contributions

Conceptualization, K.S. and D.L.; Methodology, K.S., D.L. and C.M.; Software, D.L.; Formal Analysis D.L.; Investigation, D.L.; Writing – Original Draft, D.L.; Writing – Review & Editing, K.S., D.L., and C.M.; Resources, D.L. and C.M.; Supervision, K.S.; Funding Acquisition, K.S.

## Acknowledgements

This work was supported by the Howard Hughes Medical Institute and the NINDS. We would like to thank Peri Kurshan, and Claire Richardson in the Shen lab for careful reading of this manuscript, and for their insightful suggestions. We would also like to thank Matthew Schwartz and the Jorgensen lab for providing reagents (*ox699* and the SapTrap kit).

## Methods

### C. Elegans strains and maintenance

C. Elegans strains were maintained according to standard methods, as previously described (Brenner et al. 1974). Strains were maintained at 20 degrees Celsius on plates seeded with OP50 strain E. Coli bacteria. Strains: The following *C. Elegans* strains were used in this study: TV17121 *wyIs639* [*Pdat-1::gfp::rab-3*], TV15118 *wyIs552* [*Pdat-1::unc-10::3x-gfp; Pdat-1::gfp::rab-3*], TV15123 *wyIs557* [Pdat-1::gfp::elks-1; Pdat-1::TdTomato::rab-3], TV18524 *wyIs692* [*Pdat-1::3x-gfp-novo2::syd-2; Pdat-1::mCherry*], TV15731 *wyIs552 unc-104 (e1265),* TV15727 *wyIs557 unc-104 e(1265)*, TV21685 *wyIs692 unc-104*(*e1265*), TV18112 *wyIs557 unc-104* (*wy673*), TV16174 wyIs557 *syd-2 (wy5)*, wyIs557 *syd-1 (ju82) syd-2 (wy5)*, TV16176 wyIs557 *arl-8(wy271)*, TV16175 *wyIs557 arl-8(wy271) syd-2(wy5*), TV18036 *wyIs639 unc104(wy673*), TV21853 *wyIs639 arl-8 (wy271),* TV22603 *wyIs639 syd-1 (ju82) syd-2 (wy5)*, TV21223 *ox699* [*gfp::flp-on::rab-3*]; wyEx8715 [*Pdat-1::flp*]. *elks-1(wy1162)* [*Pdat-1::flp, Pdat-1::cytoplasmic mCherry*].

### Imaging and Micrograph Generation

For all imaging experiments, described in detail below, imaging was performed using a Zeiss Observer.Z1 spinning disk confocal microscope Hamamatsu C9100-13 EM-CCD camera with a Zeiss 40X/1.3 or Zeiss 63X/1.4 objective. Images and movies were recorded using Metamorph Imaging software (Molecular Devices).

### Steady-state imaging and micrograph generation

Worms were mounted on 5% agarose pads in 2.5 mM levamisol + 0.225 mM 2,3-butanedione monoxime (Sigma) diluted in M9 buffer, and imaged up to 1 hour after sealing slide with vaseline. Metamorph Imaging software Version 7.7.9.0 (Molecular Devices) was used to acquire images. 50mW Coherent Sapphire Lasers lines of 488 and 561 nm were used. For still images, z-stack images of the worm were taken, and then stitched together using Matlab code. Axons were cropped out of the images using the “Straighten to Line” function in ImageJ. Subsequently, axons were grouped together into one image using Matlab code, and were also analyzed to generate plots of puncta intensity along the axon using Matlab code (figures 2, 4, and 7).

### Long term time-lapse imaging and kymograph generation

For Figure 3 and Supplemental Figure 7, we bathed L2-L3 worms in .4 mM levamisol diluted in M9 buffer for 30 minutes and then transferred worms individually to an M9 droplet using a glass hook. The M9 droplet was then flipped over onto a 5% agarose pad. 1 ul of .05 um microsphere beads (Cat # 08691 Polysciences Inc.) were added to immobilize the worms, and coverslip was sealed with vaseline and imaged for up to 2.5 hours after mounting. We imaged the axon outgrowth and synapse formation process by taking a z-stack frame of the PDE axon for 2.5 hours. Axons isolated using the ImageJ “Straighten to Line” function, and kymographs were subsequently generated.

### Short-term time-lapse imaging of vesicle trafficking and kymograph generation

Figures 4 and 6: Worms were mounted in 6 mM muscimol diluted in M9 buffer on 5% agarose pads in M9 buffer for 2 minutes. Then 1 ul .05 um microsphere beads (Cat # 08691 Polysciences Inc.) were added to the immobilize the worms and the slide was sealed with a coverslip and vasline. Animals were then imaged for up to one hour after initial sealing of slide. Figure 6: All strains were imaged using 100X/1.4 Zeiss objective and 1.6X opto-var magnification, and identical laser/camera settings, as set in Metamorph. Figure 4: All strains were imaged with 100X/1.4 Zeiss objective without opto-var magnification and identical camera/laser settings, as set in Metamorph. For all experiments in Figures 4 and 5, axon was first photobleached to reduce background fluorescence intensity. Photobleaching was accomplished using a 4-sec laser pulse from a UV laser line mosaic system (Andor), which allows the illumination of a targeted ROI in an image. Animals were then imaged with 488 nm laser using 100ms exposure times, at 7.5 Hz (Image was taken every 133 ms), for 60 seconds. Axons were cropped out of the images using the “Straighten to Line” function in ImageJ. Subsequently, kymographs were generated using Matlab code, and subsequently manually analyzed for transport events while blind to the genotype of the kymograph (figures 4, and 5).

### Experimental Design and Statistical Analysis

Figures 2, 4, 5, 6, and 7: One way ANOVA or Kruskal-Wallis calculations were performed using the anova1 or kruskal-wallis function in matlab. Then, the Tukey-Kramer or honestly significant difference (hsd) test was used in the multcompare function with different alpha values (a* = .05, a** = .01, a*** = .001), also in Matlab. Figure 1F: One way anova was run separately for *ox699* and *wyIs639*. Subsequent tukey’s tests showed no significant differences between bins 5-9 for either genotype. Figure 6: A kruskal-wallis test was performed on trafficking data for Figures 6E, 6F, (event frequencies) and Figures 6G, 6H (run lengths). Then Tukey-Kramer or honestly significant difference (hsd) tests were performed using Matlab’s multcompare function. Figure 4: For all panels in this figure, either a one-way ANOVA (Figures 4I-J) or Kruskal-Wallis (Figures 4K-L) were used depending on whether or not a gaussian distribution of the data was assumed. Subsequently, a Dunnet’s multiple comparisons test was used to determine significant differences between each of the mutant genotypes and the *wild-type* control. All data for this figure were analyzed using GraphPad’s PRISM 7 data analysis software. Figure 3: GFP::ELKS-1 and tdTomato::RAB-3 Co-localization: A two-way ANOVA was performed in GraphPad’s PRISM, followed by Tukey’s multiple comparisons tests to compare individual age cohorts.

