## Supplementary Materials for "Axonal transport and active zone proteins regulate volume transmitting dopaminergic synapse formation"

### Supplemental Figure 3A

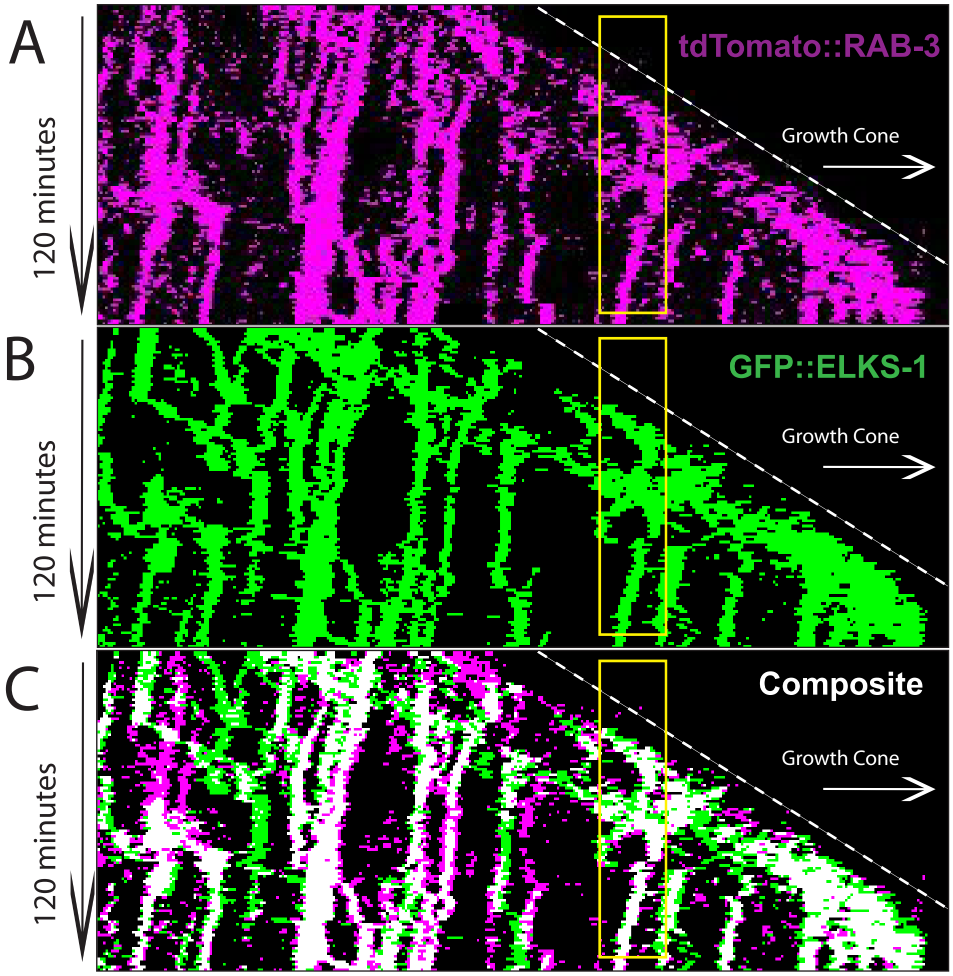

### Supplemental Figure 3B

A

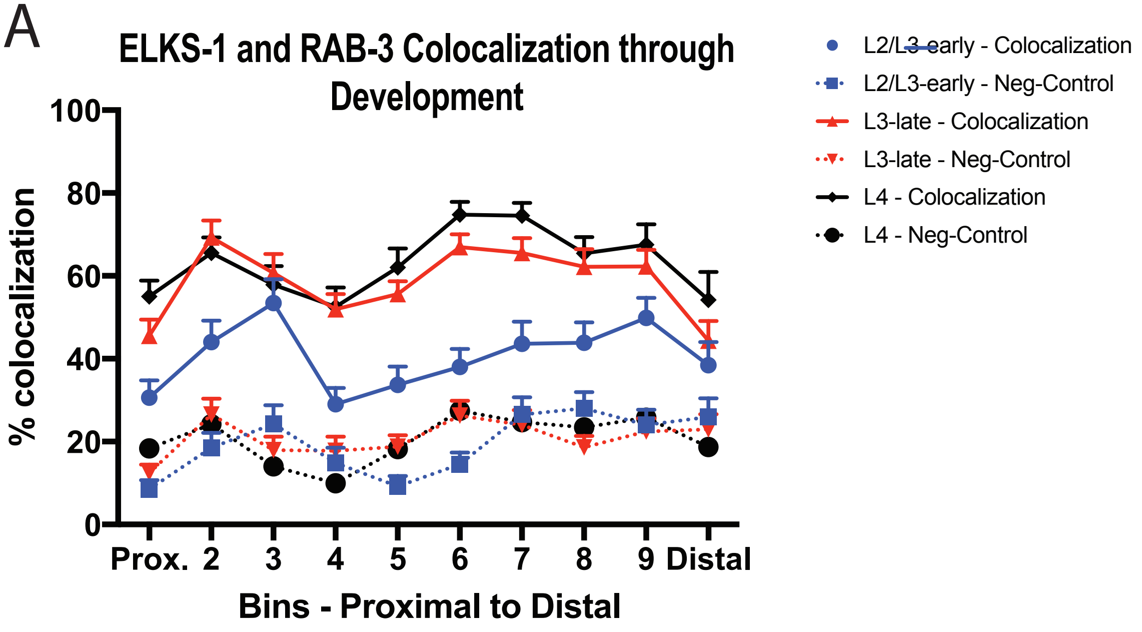

### Supplemental Figure 3C

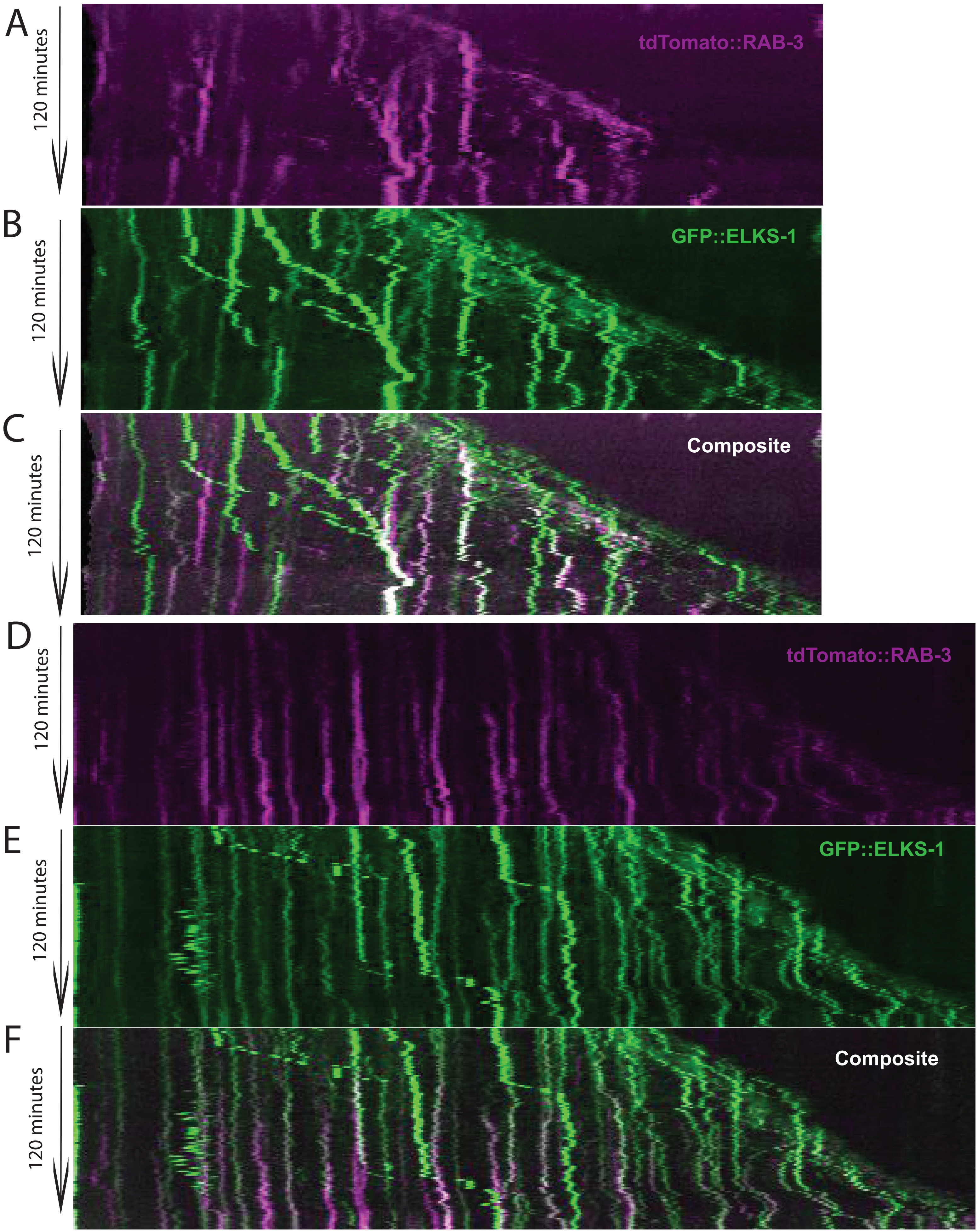

### Supplemental Figure 3D

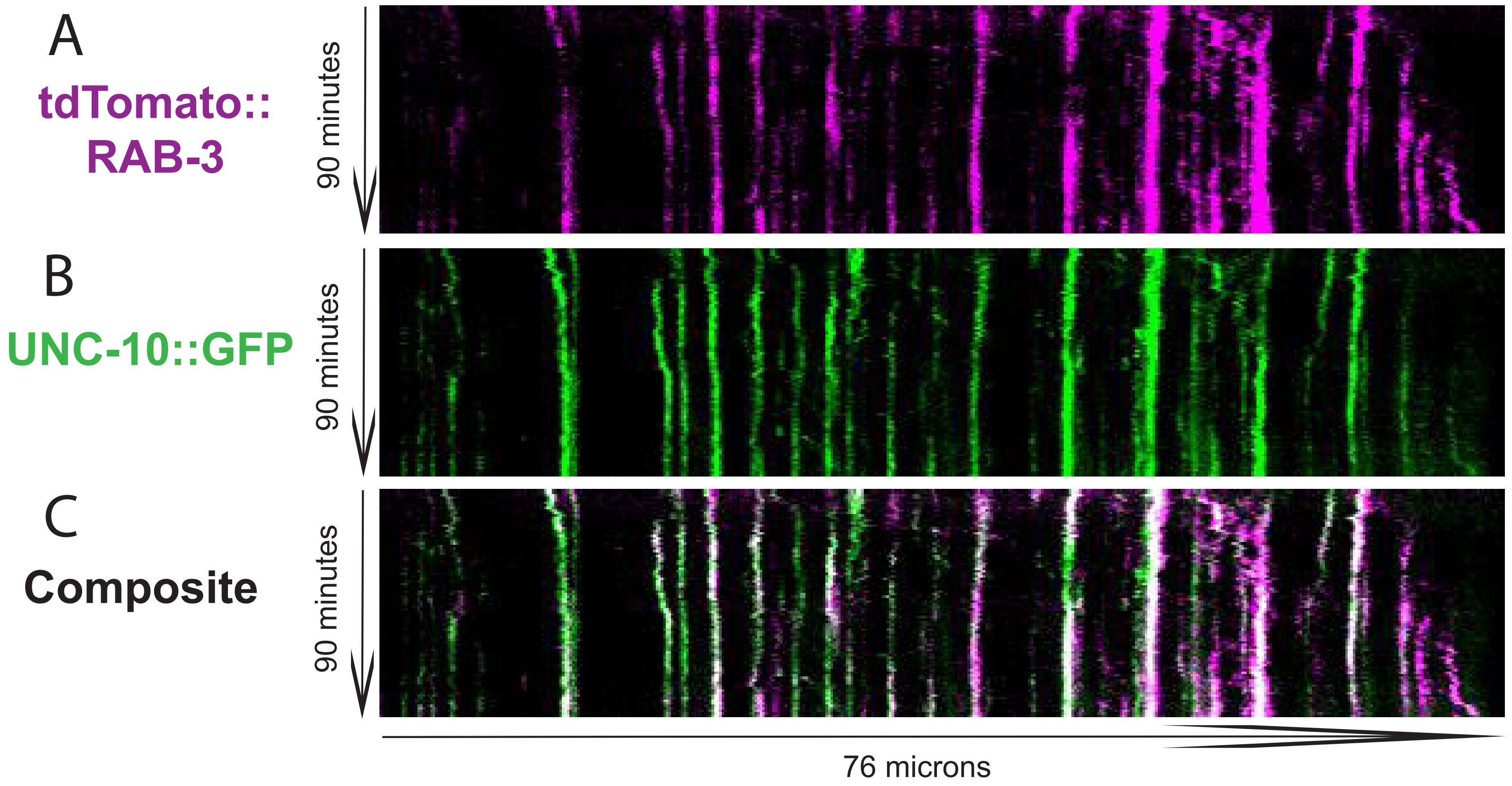

### Supplemental Figure 6

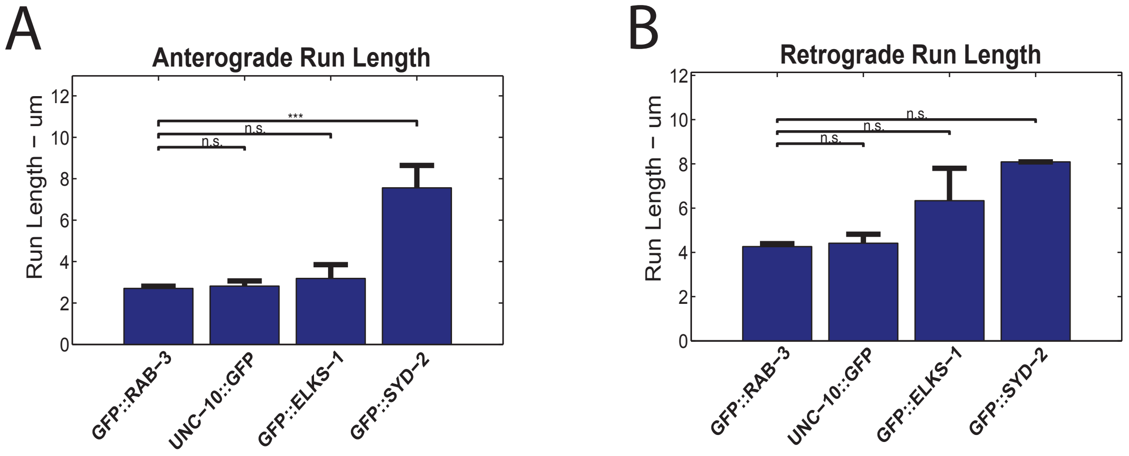

### Supplemental Figure 7

F

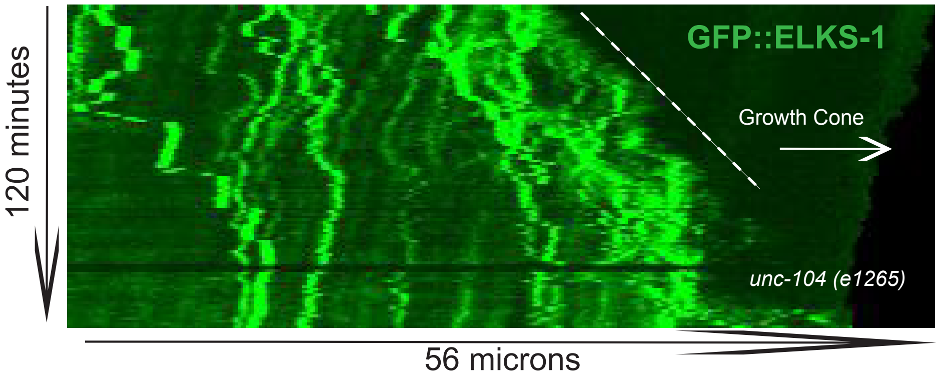
